# Dynamic network partnerships and social contagion drive cooperation

**DOI:** 10.1101/430843

**Authors:** Roslyn Dakin, T. Brandt Ryder

## Abstract

Both reciprocity and positive assortment (like with like) are predicted to promote the evolution of cooperation, yet how partners influence each other’s behavior within dynamic networks is not well understood. One way to test this question is to partition phenotypic variation into differences among individuals in the expression of cooperative behavior (the “direct effect”), and plasticity within individuals in response to the social environment (the “indirect effect”). A positive correlation between these two sources of variation, such that more cooperative individuals elicit others to cooperate, is predicted to facilitate social contagion and selection on cooperative behavior. Testing this hypothesis is challenging, however, because it requires repeated measures of behavior across a dynamic social landscape. Here, we use an automated data-logging system to quantify the behavior of 179 wire-tailed manakins, birds that form cooperative male-male coalitions, and we use multiple-membership models to test the hypothesis that dynamic network partnerships shape within-individual variation in cooperative behavior. Our results show strong positive correlations between a bird’s own sociality and his estimated effect on his partners, consistent with the hypothesis that cooperation begets cooperation. These findings support the hypothesis that social contagion can facilitate selection for cooperative behavior within social networks.

## INTRODUCTION

Cooperation is an emergent property of social interactions within a network, yet our understanding of how cooperative behavior emerges and is maintained is still major problem in evolutionary biology. Two processes that favor the evolution of cooperation are reciprocity, wherein cooperative behavior is socially contagious, and positive assortment, wherein non-random structured interactions create clusters of cooperators in the network [1–4]. These processes are non-exclusive, because any source of reciprocity that causes individuals to act more like their cooperative neighbors will also contribute to the process of positive assortment. Hence, a major question is how these social processes within complex networks influence variation in cooperative behavior and the emergence of sociality.

Answering this question requires partitioning phenotypic variation in cooperation into two major sources (Fig. 1a): (1) differences among individuals in the expression of a behavior due to their intrinsic biology (the “direct effect”), as well as (2) plasticity within individuals induced by the social environment (the “indirect effect”). This approach, largely derived from quantitative genetics (i.e., interacting phenotypes and indirect genetic effects [5–8]), accounts for the multilevel nature of behavior and can be used to quantify social influence while accounting for other sources of variation [9,10]. A key feature of this framework is its focus on repeatable individual differences [9]. This is important because repeatable phenotypic variation is the raw material upon which selection acts [11].

**Fig. 1.**
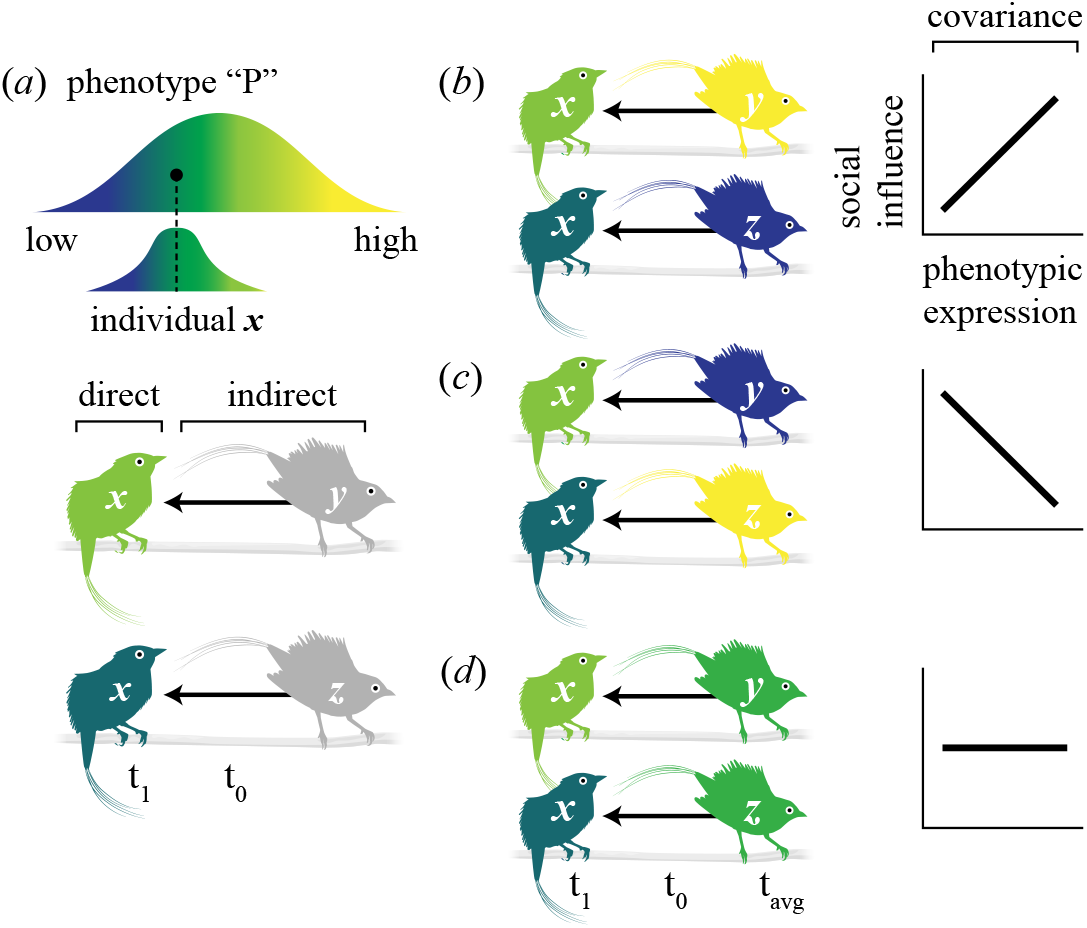
A conceptual framework for social influence. (a) A phenotype “P” varies within and among individuals. In this example, yellow indicates high expression of the trait whereas blue indicates low expression of the trait. The upper histogram shows the hypothetical phenotypic distribution for a population, whereas the lower histogram represents the narrower range of phenotypic values expressed by a given individual, X, within that population. Individual phenotypes are the result of both intrinsic biology (the “direct effect”) and plasticity induced by the social environment (e.g., the “indirect effect” as X interacts with conspecifics Y and Z). This indirect effect represents how interactions with Y and Z at an earlier time, t_0_, can differentially influence X’s phenotype at time t_1_. Note that X would also influence Y and Z’s phenotype at time t_1_ in the analysis, but for simplicity, we show only one direction of social influence here. (b-d) Because Y and Z also express phenotype P, there may be covariance between an individual’s average phenotypic expression (t_avg_) and his average social influence. For example, (b) Y and Z’s average phenotypic expression may be positively correlated with their social influence on X’s phenotype. This positive covariance is predicted to generate social contagion. Alternatively, phenotypic expression may be (d) unrelated or (c) negatively related to social influence, the latter of which would generate social inhibition.

Variance partitioning can also be used to quantify among-individual variation in social influence (i.e., the extent to which individuals differentially alter the expression of their partner’s phenotype). This is important given growing evidence that the social environment can alter phenotypic expression and subsequent selection dynamics (reviewed in [9,10]). Moreover, estimates of social influence as a trait are key to understanding how behaviors are transmitted or inhibited within social networks [12]. When an individual’s average phenotypic expression positively covaries with his social influence (e.g., the most cooperative individuals elicit their partners to cooperate, Fig. 1b), behaviors can be transmitted via social contagion among partners within a network [12]. This social contagion is hypothesized to promote the spread of cooperative behavior and facilitate selection for cooperation [7, 13–15]. Conversely, negative covariance between an individual’s average expression of a trait and his social influence (Fig. 1c) creates social inhibition which prevents the spread of, and constrains selection for, a specific behavioral trait within the network [16–18]. Finally, there may be no relationship between average expression and social influence among interacting phenotypes (Fig. 1d).

These theoretical principles highlight how social influence can promote and/or constrain the emergence of population-level processes such as cooperation [10,19,20], but empirical tests have been limited. Recent experiments have demonstrated that cooperative and social behaviors can be transmitted via social contagion and dynamic network partnerships in flies, fish, and humans [12,14,15,21–23]. To date, however, these laboratory studies have considered simple dyadic or small group interactions that do not reflect realistic social complexity [but see 24], and may overestimate social influence [8,25]. Thus, a complete understanding of social influence will require collecting repeated measures of behavior while accounting for who interacts with whom, and the frequency with which these interactions occur in a dynamic social network.

Here, we address this gap by testing how the social environment influences cooperative behavior in a lek-breeding bird, the wire-tailed manakin (*Pipra filicauda*). The social system of this species is well-studied and consists of two male status classes defined as territorial and non-territorial (or ‘floater’) males [26]. Both status classes engage in repeated social interactions in the context of male-male display coalitions which function both in the maintenance of dominance hierarchies and in attracting females [27; see also Movie 1]. At any given time, a male will have multiple coalition partners and these heterogeneous partnerships create complex reticulate social networks [28]. Moreover, although manakin coalitions are relatively stable on an annual basis [27], partnerships are dynamic on shorter daily or weekly timescales (see ESM 1) and males show substantial variation in cooperation [28,29]. Previous work on wire-tailed manakins has established that cooperative behavior within the social network has clear fitness benefits: territorial males with more coalition partners have greater reproductive success [30], and floaters with more coalition partners have an increased probability of territorial inheritance [28]. Attaining coalition partners is thus essential for floater fitness because territoriality is a prerequisite for access to mating and reproductive success [30].

To evaluate the influence of social network interactions (i.e., coalition partnerships) on social behavior, we leveraged our knowledge of wire-tailed manakin behavior to develop an autonomous proximity data-logging system (Fig. 2). The proximity approach allowed us to obtain repeated measures of behavioral phenotype for a population of 179 males and to characterize the identity and frequency of cooperative coalition partnerships within the social network. We chose four social behaviors that are expected to promote reproductive success and thus be under selection. “Effort” is a measure of daily attendance on territories where the coalitions are formed and is a behavior known predict male mating success in other lekking taxa [31]. “Degree” and “strength” measure the number of cooperative coalition partners and the frequency of those partnerships within the network, respectively. These measures of cooperation are known to affect social ascension and reproductive success in manakins and other organisms [28,30,32]. Social “importance” measures a male’s position within the broader social network by characterizing the exclusivity of his coalition partnerships. Exclusivity is contingent on the proportion of direct and indirect social relationships (i.e., how frequently your partner displays with you vs. with his other coalition partners); recent work suggests that these indirect network connections often capture among-individual differences in social behavior [33]. Moreover, in manakins, the exclusivity and subsequent stability of coalition partnerships can increase elements of display coordination and signal intensity, both of which are preferred by females [27,34].

**Fig. 2.**
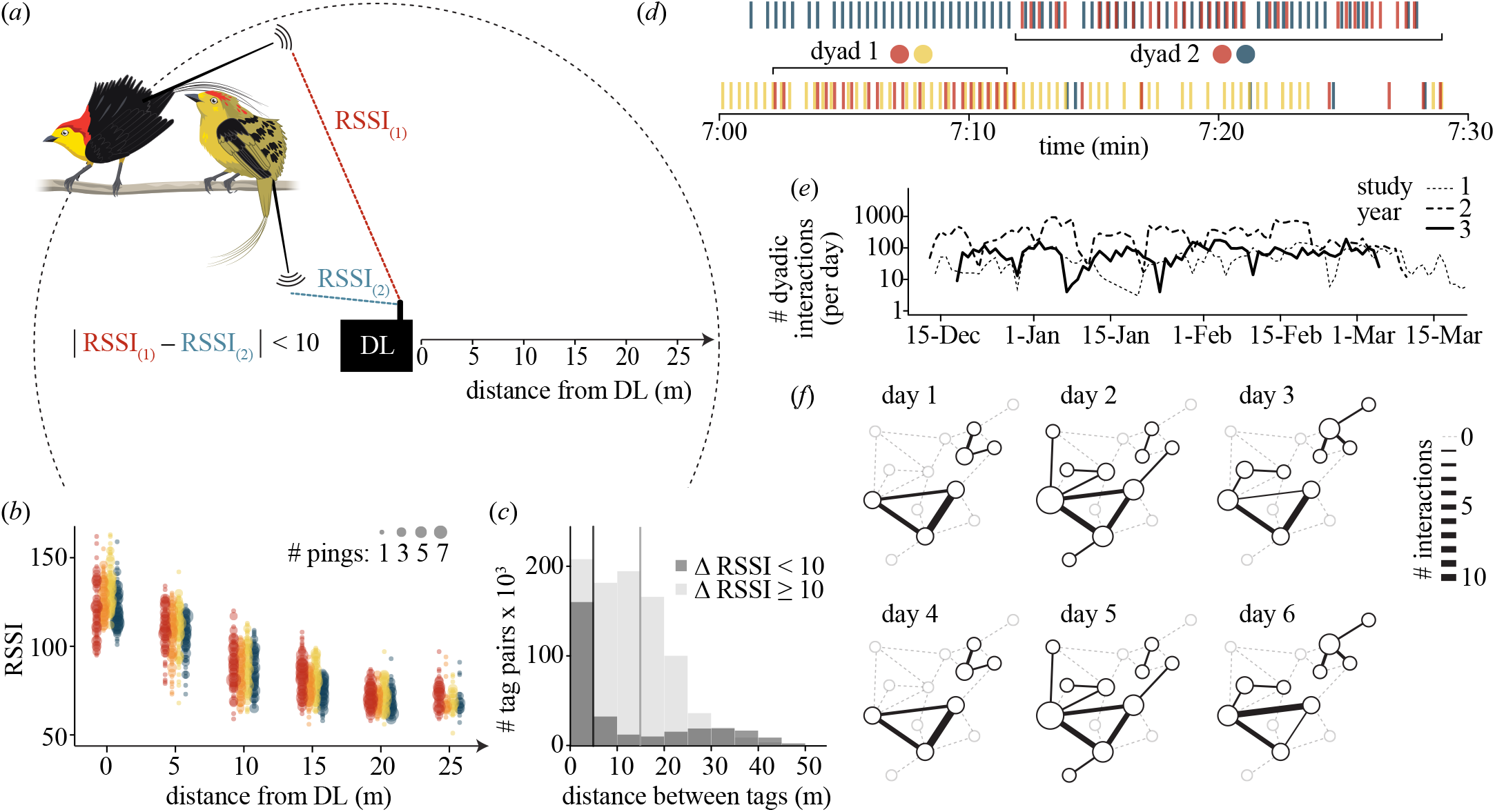
Social interactions were tracked using an automated proximity data-logging system. (a) Each bird was tagged with a small transmitter that broadcasts a unique digital code at 20 s intervals. Tag pings were received by stationary data loggers (DLs) set with a fixed detection radius to encompass a male’s territory where cooperative displays take place. (b) The tag signal strength, RSSI, decays with distance from the DL. Hence, we define an interacting dyad using a conservative spatiotemporal threshold wherein individuals had to be detected within 45 s of each other and with ΔRSSI < 10. (c) A ground-truthing analysis demonstrated that tag pairs that meet this threshold are typically ≤ 5 m apart (vertical line = median), corresponding to a range that would permit both visual and acoustic contact. For comparison, events with ΔRSSI ≥ 10 have a greater median inter-tag distance of 15 m. The data in (b) and (c) are based on the same ground-truthing experiment. (d) An example of detection data for DLs at two neighboring territories. The resulting time series illustrates two interactions, dyad 1 and dyad 2, that are delimited by horizontal brackets. Note that birds occasionally ping the same DL around the same time without meeting the ΔRSSI spatial threshold. (e) Overall, this automated system recorded approx. 37,000 social interactions among 179 birds. (f) An example of daily interactions within a subset of the population social network illustrating the dynamic nature of the study system. Each node in the network represents an individual male sized according to degree. Data are shown for 14 of the males recorded from January 1-6 in 2016, with solid edges representing interactions that occurred on a given day. Dotted edges are interactions that did occur but not on that day.

Here, we use multiple membership models to evaluate the extent to which an individual’s intrinsic biology (direct effect) and his repeated social network interactions (indirect effect) can explain variation in cooperative behavior. Our analysis had three aims. First, we estimated the direct effect as the proportion of total phenotypic variance attributed to differences among individuals (i.e., the repeatable component of X’s own behavior in Fig. 1a). Second, we estimated the indirect effect as strength of social influence on phenotypic variation within individuals (i.e., the repeatable component of Y and Z’s influence on other individuals in Fig 1a). Our analysis also captures the reciprocal nature of social influence in each coalition partnership. A significant indirect effect in this analytical framework has two main implications: it tests how much of the total phenotypic variance can be explained by social influence, but also, it measures how individuals vary in their social influence within the network by accounting for the frequency of interactions between coalition partners. Third, we examined the transmission of behaviors among coalition partners by testing for social contagion (Fig. 1b-d). Given that social contagion is defined as the spread of a particular behavior among partners within a social network [12], we estimated the correlation between an individual’s average expression of a trait and his social influence on that same trait in his coalition partners. A positive covariance between these two repeatable components of variation is consistent with the hypothesis that the expression of a given behavior is socially contagious. We compared all of these results to null models of the network data in which the direct effects and indirect effects had been removed by permuting individual ID labels.

## MATERIAL AND METHODS

### Study System and Field Methods

We studied a color-banded population of male wire-tailed manakins (*P. filicauda*) at the Tiputini Biodiversity Station, Orellana Province, Ecuador (0° 38’ S, 76° 08’ W, approx. 200 m elevation) across three breeding seasons (November to March 2015-18; hereafter years ‘15-16’, ‘16-17’, and ‘17-18’). Male status (i.e., territorial or floater) was determined by direct observation in the field following Ryder et al. [28].

To collect repeated measures of the behavior and social environment of male manakins, we developed an automated proximity system shown in Fig. 2 [35]. Each individual was outfitted with a coded nano-tag (NTQB-2, Lotek Wireless; n_15-16_ = 100 males that were tagged and color-banded, n_16-17_ =114, n_17-18_ = 82; and n_total_ = 179). Annual sample sizes are non-exclusive because many males were tagged in multiple years (see details below). The sample size was not pre-determined, but instead the aim was to capture all individuals and monitor as many social network interactions as possible. Proximity data-loggers (hereafter DL; SRX-DL800, Lotek Wireless) were placed within the lek territories where the manakins engage in their cooperative displays (Fig. 2a; see ESM 1). DLs recorded continuously for 10 hours from 0600 to 1600, such that each day constitutes a repeated measure of behavior. DLs were deployed at a given lek for an average of ~6 days (± 1 SD) per sampling interval and moved among leks on a rotating schedule. In total, we recorded 29,760 hours of proximity data over 249 days (15-16: 49 territories, mean = 16 recording days per territory; 16-17: 54 territories, mean = 21 days; 17-18: 48 territories, mean = 22 days, SD = ±4 days in all years).

### Data Processing

To define distinct social interactions from the proximity data, we used a rule set with spatial and temporal thresholds illustrated in Fig. 2a-d. This rule set was based on 16 years of previous study demonstrating that spatial co-occurrence within a manakin territory is a both a necessary prerequisite for, and an accurate predictor of, cooperative display interactions among males [28,35]. We also used a ground-truthing experiment to confirm that the spatial threshold would identify birds in close proximity (Fig. 2b-c; see ESM 1).

We used the proximity data to calculate repeated measures of four behavioral phenotypes defined on a daily basis. (1) Effort is the number of unique DL pings by a focal individual. (2) Strength is the sum of a male’s edge weights within the network. (3) Degree is the number of unique edges or direct links a male has within the network. Strength and degree are thus measures of the frequency of cooperative behavior and the number of cooperative coalition partners, respectively. (4) Importance is a measure of the exclusivity of a male’s coalition partners, on a scale from 0 to 1. To calculate importance, we first found the proportion of each partner’s interactions that were with a focal male. Then, we took a weighted average using the focal male’s interaction frequencies. Thus, a male whose partners often interact with other individuals would obtain a low score for importance, whereas a male whose partners interact with him exclusively would obtain a score of 1. Importance is thus akin to an indirect network metric that accounts for both direct and indirect social interactions (reviewed in [33]). Additional details, descriptive statistics for the behavioral phenotypes, and their correlations are provided in ESM 1 (see Fig. S1 and Tables S1, S2).

### Statistical Analysis

#### Assortment

To quantify network assortment, we calculated the assortativity coefficient, ρ, for each of the four phenotypes. This analysis was based on a single average value for each individual and a single social network compiled over all three years of study. We first derived Newman’s assortativity coefficients using the igraph package 1.1.2 in R [36,37], and verified that the results were identical to weighted network assortment coefficients for continuous phenotypes using the assortnet package 0.12 [38,39]. To test whether phenotypic assortment was statistically significant, we determined the 95% confidence interval using the jackknife resampling method.

#### Direct and Indirect Effects

We used multiple-membership models (hereafter, MMMs) for partitioning the phenotypic variance between the direct and indirect effects. MMMs are multi-level models that incorporate a heterogeneous weighted random effect [40], making it possible to capture variation in dynamic network partnerships. The key advantage is that MMM analyses can be weighted according to the frequency with which different social relationships occurred.

We fit MMMs using the brms package 2.5.0 in R [37,41] and the model syntax shown in Box 1. The response variable was an individual’s behavioral phenotype on a particular day (t_1_); we also included the bird’s identity as a random effect, as well as his top four most frequent partners on the previous day (t_0_) in the multiple-membership structure, weighted by their interaction frequencies. Thus, our analysis asks, how is a bird’s behavior influenced by his own identity and who he interacted with on the previous day? Although a few birds had more than four partners in a single day, we limited our analysis to the top four because additional partners comprised less than 2% of a bird’s daily total in our data. When an individual had fewer than 4 partners on a given day, we distributed one, two, or three of his realized partner IDs and their interaction frequencies over the remaining empty slot(s), while maintaining each partner’s total weight as a constant. This procedure does not influence the results because MMM analysis normalizes the weights within the focal individuals [40].

We also included fixed effects of focal status (territorial vs. floater), study year, and an individual’s number of years of tenure in the present study, as well as random effects of date and lek in all models. By including year, date, and lek, our estimates of the direct and indirect effects account for spatial and temporal variation in the shared non-social environment that may influence focal and partner behavior (e.g., [42–44], Table S1). Years of tenure was included instead of age, because age was not known for most (69%) of the social interactions. Hence, our analysis computes the proportion of phenotypic variance due to direct and indirect effects after accounting for status and any spatial/temporal dynamics. Although feedback loops are implied [8], we do not consider the influence of second- or higher-order connections among focal and partner phenotypes [45,46].

Prior to modelling, three of the four behavioral phenotypes were log-transformed (effort, strength, and degree) so that Gaussian model assumptions were met, and all phenotypes were standardized (mean 0, SD 1). We used the default uninformative priors and stored 2,000 samples from each of four independently-seeded chains for each phenotype, verifying that the convergence statistics in the model output were all equal to 1. We then obtained estimates of the variance components for each phenotype by pooling the posterior values from all four chains. Posterior distributions for the direct and indirect effects were calculated as *Var_focal_*/*Var_total_* and *Var_social_*/*Var_total_*, respectively [9], wherein *Var_focal_* is the variance due to the identity of the focal individuals; *Var_social_* is the variance due to the MMM structure (i.e., the social environment); and *Var_total_* is the sum of all variance components (including focal, social, date, lek, and the residual/unexplained variance). Our sample size for this analysis was 2,935 daily measures of 144 focal individuals (see ESM for details). Note that this is smaller than the number of tagged birds, because a focal bird had to have a known social environment on the previous day to be included in the MMM analysis. Of these 144 males, 69 were tracked in one year, 49 in two years, and 26 in all three study years; 8 floater individuals also attained territorial status during the study.

###### Box 1. Syntax for weighted MMM analysis of direct and indirect effects in brms

scale(Focal.phenotype) ~ focal.status + study year + tenure, random + (1|Focal.ID) + (1|mm(Partner.ID1, Partner.ID2, Partner.ID3, Partner.ID4), weights=cbind(frequency.1, frequency.2, frequency.3, frequency.4)) + (1|date) + (1|lek) Box 1. Syntax for weighted MMM analysis of direct and indirect effects in brms scale(Focal.phenotype) ~ focal.status + study year + tenure, random + (1|Focal.ID) + (1|mm(Partner.ID1, Partner.ID2, Partner.ID3, Partner.ID4), weights=cbind(frequency.1, frequency.2, frequency.3, frequency.4)) + (1|date) + (1|lek)

#### Null Models

Given the non-independence of network data, we used a null model framework to compare our posterior estimates with the same analyses performed on randomly permuted data [47,48]. The null permutations preserved all structure in the data but permuted the ID labels, which were swapped among individuals in the same daily recording session. This is equivalent to a node-label network permutation. Given that territorial and floater birds differ [35] (Table S1), the permutation was also constrained to swap ID labels among status-matched individuals recorded on the same day. Hence, these null datasets (n =1,000) preserved spatiotemporal sources of variance/covariance, but disrupted sources of variance/covariance that could be attributed specifically to identity. Null MMMs were fit using the same syntax presented in Box 1 and thus provided a strong test of the identity-based effects that are the focus of our analysis (see ESM 1 for details).

#### Contagion

Social contagion generates a positive covariance between a bird’s own phenotypic expression and his social influence on others’ expression of the same trait (Fig. 1b-d). To determine this covariance, we used the posterior estimates of phenotypic expression and social influence for each male in the population from the fitted MMMs. We used the posterior medians to test the correlation between an individual’s own phenotypic expression and his social influence on each of the four behavioral phenotypes. That is, do individuals that express consistently high levels of a phenotype also elicit that same phenotype in others? To verify that these associations were significant, we compared the observed correlation statistic (r) with that derived from the null permutations, which preserved spatiotemporal sources of covariance to provide a conservative test. Finally, because directly estimating covariance from the MMMs is not currently possible ([41,49]; Bürkner and Hadfield, pers. comm.), we also performed a sensitivity analysis to validate this approach (ESM 1). The sensitivity analysis demonstrated that in models with a large sample size like ours, the bias in the direct and indirect effects caused by covariance is very small, and much smaller than the uncertainty inherent in these estimates (Fig. S2). Thus, any bias is not expected to affect our conclusions.

## RESULTS

Three of the four phenotypes, strength, degree, and importance, were positively assorted in the social network (“like with like”; strength ρ = 0.24 ± SE 0.06; degree ρ = 0.33 ± 0.05; importance ρ = 0.27 ± 0.05), whereas the network assortment was not significantly different from 0 for effort (ρ –0.01 ± 0.06). All four phenotypes also had statistically significant individual differences in expression, such that the direct effects could explain ~12-30% of the total variance (Fig. 3a, Table S3, all p < 0.006). Because these percentages are moderate, this result also demonstrates substantial within-individual plasticity in all four behavioral phenotypes.

**Fig. 3.**
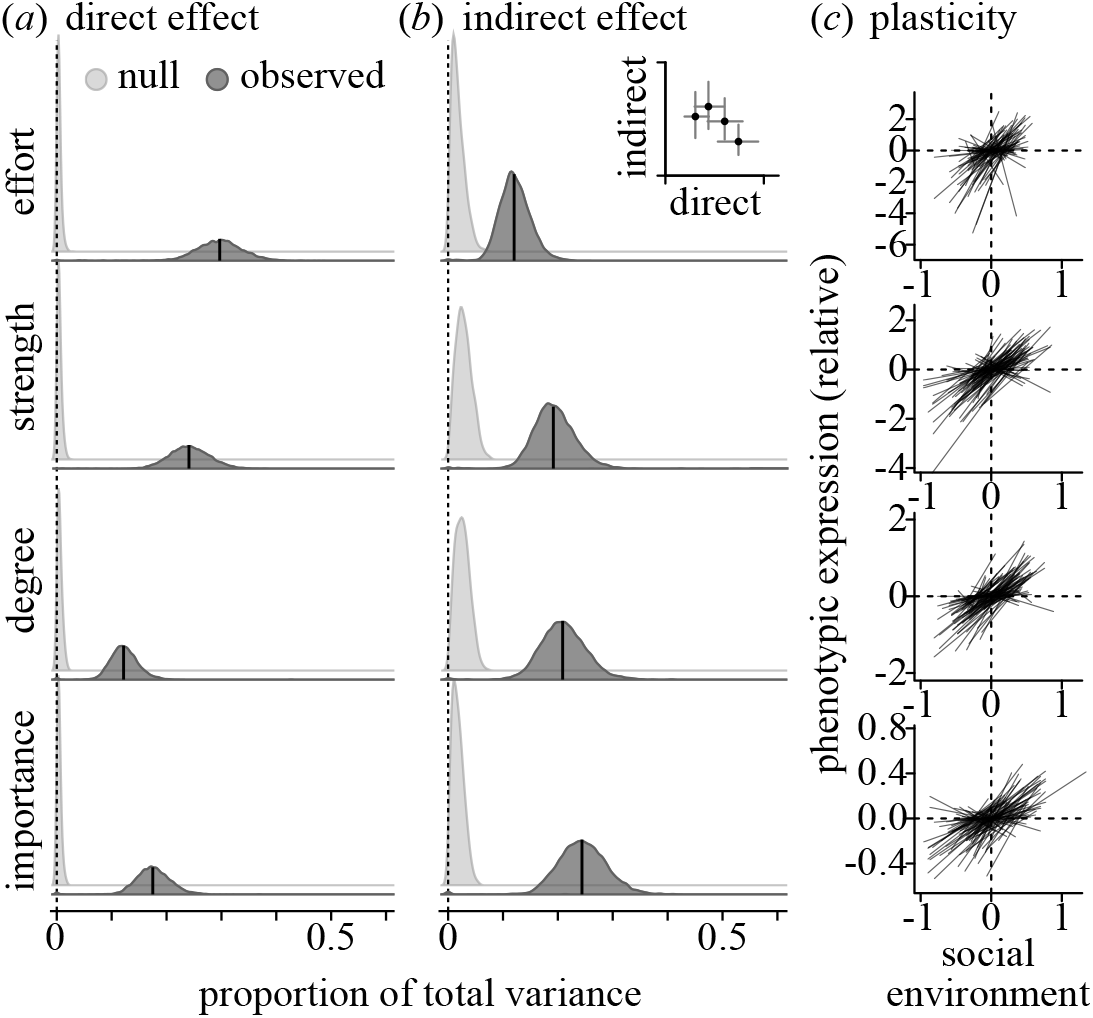
Manakins differ both in their social behavior and their influence on others. (a-b) The proportion of total phenotypic variance in four behavioral phenotypes attributed to (a) individual differences (the direct effect) and (b) social influence (the indirect effect), as defined based on partners from the previous day. For each phenotype, the observed result is significantly greater than the null expectation. The vertical lines show the posterior median. Phenotypes with larger direct effects tend to have weaker social influence (inset, b). The axes on the inset span 0 to 0.4. Note that the tails of all distributions are extended beyond the range of the data for visual continuity. (c) Within individuals, phenotypic expression depends on the previous day’s social environment. On the y-axes, individual phenotypic expression values are centered to show change relative to an individual’s mean expression; values for effort, strength, and degree were also log-transformed prior to centering. The x-axis represents the average influence of a bird’s partners on the previous day. A separate linear fit is shown for each focal bird observed on at least 10 days (n = 95). See Tables S3 and S4 for detailed results.

We next asked if the social environment could explain some of this within-individual variation in behavior. We found that between 12-24% of the total variance could be attributed to indirect effects caused by the dynamic network interactions on the previous day (Fig. 3b, Table S3, all p < 0.005). The phenotypes with the larger direct effects (effort and strength) also tended to have smaller indirect effects, and vice versa (inset, Fig. 3b). These significant indirect effects also imply that social influence is itself a trait that varies among individual male manakins. Moreover, our null models demonstrate that indirect effects of this magnitude do not arise by chance in permuted data with the exact same phenotypes and network topologies.

To further visualize these indirect effects, we also plotted within-individual plasticity in phenotypic expression in relation to the social environment, as defined by average partner social influence from the previous day (Fig. 3c). These plots show how individuals adjust their behavior based on their dynamic social environment on the previous day.

Finally, we asked if indirect effects as our measure of social influence could drive the social contagion of behavior. We found strong positive correlations between expression and social influence for three of the four phenotypes: strength, degree, and importance (Fig. 4), all of which also exhibited positive assortment. Moreover, these estimates of covariance were greater than expected in our permuted data (Fig. S3). These results suggest that individual manakins with consistently high strength, degree, and importance tend to elicit increases in those respective phenotypes in their partners. In contrast, although greater effort was also elicited by particular individuals (Fig. 3), its lack of a significant positive correlation suggests that effort is not socially contagious (Fig. 4).

**Fig. 4.**
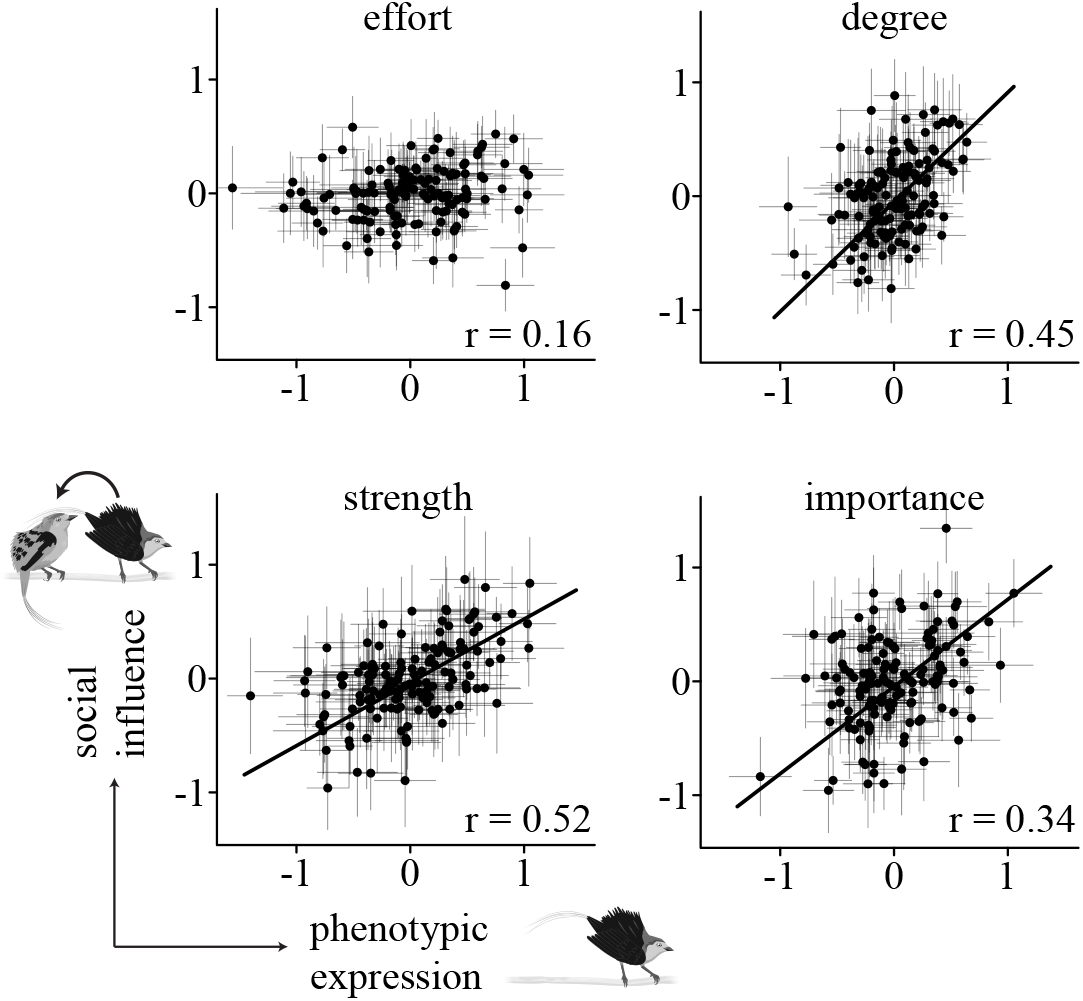
Positive covariance indicates that cooperative social behaviors are contagious. Each graph shows an individual’s estimated social influence (y-axis) in relation to his own phenotypic expression (x-axis) for that same behavior (n = 142 individuals analyzed as both focal and partner). Strength, degree, and importance all have strong positive correlations that are significantly greater than 0 and greater than the null expectation from the permuted data, indicating that indirect effects can facilitate the spread of these behaviors through the social network. In contrast, the correlation for effort is not statistically significant. Data points and error bars are derived from the posterior medians and standard deviations, respectively. The regression fit lines account for posterior estimates of the error in both variables. Pearson’s correlation coefficients are also given on the lower right of each panel.

## DISCUSSION

Social contagion facilitates the evolution of positive assortment and cooperation [4,12], yet understanding how dynamic network interactions shape variation in cooperative behavior has remained challenging outside of the laboratory. Here, we monitored a large population of cooperative wire-tailed manakins, and we applied a multiple-membership analysis to partition direct and indirect sources of variation in four behavioral phenotypes. We found that cooperative behavior was socially contagious: those individuals who had on average more cooperative coalition partnerships (degree) and higher frequency of interactions (strength) tended to elicit greater levels of those behaviors in their partners. Moreover, individuals who exhibited more exclusive and stable partnerships (high importance) elicited their partners to have more exclusive coalition partnerships as well. These results are notable because social connectivity as measured by the number of coalition partners has a direct link to reproductive success in this population and other species in the same family [28–30]. In contrast, we find no evidence that a male’s effort in attending the leks was socially contagious (although it was influenced by partner identity). Viewed cumulatively, our work demonstrates how multiple sources of phenotypic variation can be parsed in a network to quantify the role of social influence on behavioral plasticity in the wild. Furthermore, we demonstrate that a given behavior can be horizontally transmitted through the network via social contagion, and that this process can contribute to positive network assortment reinforcing cooperative behavior.

Over the last five years, social network analysis has rapidly advanced our understanding of the causes and consequences of social behavior [reviewed in 50]. For example, key studies have characterized individual differences in network position [51,52], patterns of phenotypic assortment [53,54], and the fitness consequences of repeated social interactions [29,30,55]. Only recently, however, have we begun to integrate network theory into our thinking about selection dynamics and how structured interactions modulate phenotypic variance [19,20]. Here, we continue to advance that theme with the first empirical evidence that dynamic network interactions and social influence can have profound effects on variation in cooperative behavior. We expect that these social processes also contribute to the emergence and maintenance of cooperation in other animal systems [10], yet empirical validation is needed.

If the observed correlation between the expression of cooperation (strength and degree) and social influence in wire-tailed manakins has some genetic basis (i.e., heritability), then our results imply that indirect effects can enhance the response to selection on cooperative behavior [5,7,13]. This is because selection acting on the genetic architecture for cooperative behavior would also facilitate a social environment that reinforces the benefits of cooperation [4]. The genetic basis of social influence could be tested in future studies that build on our approach by using genotyping to resolve the population pedigree. This will require analytical methods that can account for genetic structure in the social environment [16,18,24].

Biologists have long recognized that no phenotype is expressed in a vacuum. It is important to note that in natural settings, the non-social environment is also variable, and thus interacting individuals may share features of the environment that can be mutually influential. Although this possibility cannot be excluded here or in any observational study, our analyses account for sources of shared annual, daily, and spatial variation. Hence, the results are expected to be independent of factors that vary in space and time, such as resource availability, female activity, and climatic variation [42–44].

Another important point is that in natural settings, individuals have the capacity to choose their coalition partners and these decisions can directly impact the costs and benefits of engaging in cooperative acts [21,22,56]. For example, young wire-tailed manakins are known to increase coalition stability with age as they ascend the queue to territorial status [27]. The long-term nature of male social partnerships and behavioral plasticity in response to the dynamic social landscape [i.e., social competence, *sensu* 57] both likely facilitate the choice of similar partners (homophily) in this study system. Furthermore, homophily of socially contagious traits may further amplify or accelerate their transmission within well connected social networks [20]. Determining the relative contribution of these two processes (contagion and homophily) in a natural social network system remains a major challenge for future work and may require social transplant experiments to resolve.

Finally, our results raise several key questions for further work. First, what are the sensory mechanisms of social contagion for cooperative behavior (e.g., calls, odors, and/or gestural displays)? Second, our analysis here considered each behavioral phenotype separately, but in reality, these behaviors are not all independent (Table S2). Thus, the causal pathways for social transmission may be complex. Third, what explains the persistence of individual differences in social influence [20]? Fourth, what are the underlying physiological and mechanistic processes, and how do these mechanisms facilitate or constrain social contagion as a form of cooperative reciprocity? Work examining the hormone signaling pathways that control cooperative behavior will further elucidate the links between genotype, phenotype, and partner interactions [58]. Given that repeated heterogeneous interactions underlie virtually all animal social systems, our study provides a framework that can be broadly applied to test how social influence changes in response to physiological manipulations. Ultimately, understanding how selection shapes interactive phenotypes like cooperation will require integrative approaches that consider both mechanism and social context.

## ETHICS

All methods were approved by the Smithsonian ACUC (protocols #12-23, 14-25, and 17-11) and the Ecuadorean Ministry of the Environment (MAE-DNB-CM-2015-0008).

## DATA ACCESSIBILITY

Data are available at: https://figshare.com/s/e73a3cb48e4714898d55 [59]

## AUTHORS’ CONTRIBUTIONS

TBR and RD designed the study. TBR collected the data. RD and TBR analyzed the data and wrote the manuscript.

## ACKNOWLEDGEMENTS

We thank Ignacio Moore, Ben Vernasco, and Camilo Alfonso for their tireless effort in project organization and collecting field data. David and Consuelo Romo, Kelly Swing, Diego Mosquera and Gabriela Vinueza provided amazing logistical support at Tiputini Biodiversity Station. We thank Brian S. Evans for his brilliance in data architecture and the development of R functions. Brent Horton, Mike Webster, Joel McGlothlin, Vincent Careau’s Functional Ecology Lab at UOttawa, Niels Dingemanse, and one anonymous reviewer provided valuable comments on an earlier version of this manuscript.

